# Elevation shapes alpine snow algal blooms and their influence on albedo reduction

**DOI:** 10.64898/2026.05.12.724566

**Authors:** Pablo Almela, Scott Hotaling, J. Joseph Giersch, Helena C. L. Klip, James J. Elser, Trinity L. Hamilton

## Abstract

Snow algae darken snowpacks and accelerate melt worldwide. Although elevation strongly structures the physical conditions of mountain snowfields, its influence on snow algal traits and their effects on snowpack reflectance remains unclear. Here, we investigated snow algal composition, cellular traits, and optical properties in summer blooms across an elevational range of 1,059–3,423 m a.s.l. in the western United States, spanning two elevational gradients in the Cascade Range (CA, OR, WA) and the Rocky Mountains (UT, WY, MT). Across all samples (n = 294), snow albedo declined strongly with increasing algal cell density, indicating that total biomass, rather than pigment composition, is the dominant driver of albedo reduction. However, within Sanguina-dominated blooms (117 of 206 samples bloom samples identified across the dataset), neither relative abundance nor algal cell density varied systematically with elevation. Instead, mean cell size increased with elevation, while per-cell pigment concentrations declined, leading to higher astaxanthin:chlorophyll-a ratios driven primarily by reductions in chlorophyll-a per cell. These elevation-dependent shifts in cell size and pigment balance were consistent across both mountain ranges, indicating phenotypic acclimation to increasing environmental stress with elevation. Together, these findings link cellular-scale acclimation of a widespread snow alga to radiative processes shaping mountain snowpacks.

## Introduction

Snow cover strongly influences land–atmosphere energy exchange, primarily through the albedo–temperature feedback, in which changes in snow albedo regulate surface energy absorption and temperature (Barry et al., 2002). Although fresh snow reflects up to 90% of incoming solar radiation, albedo declines with snow metamorphism (e.g., changes in snow grain size) and with accumulation of abiotic (e.g., dust) or biotic (e.g., algae) particles that accelerate melt (Skiles et al., 2018a). Snowfields that persist into spring and summer can provide habitat for pigmented unicellular snow algae (Hoham & Remias, 2020). These snow algal blooms can be red, green, orange, or yellow depending on the dominant pigments, which vary with species composition (Almela et al., 2025a) and cell stage (Leya, 2013). Red blooms, typically dominated by *Sanguina nivaloides* (Procházková et al., 2019a; Remias et al., 2023), are the most widespread in alpine environments. Their coloration arises from high intracellular concentrations of astaxanthin, a secondary carotenoid, and its fatty-acid esters (Müller et al., 1998; Gorton et al., 2001). Carotenoid accumulation provides protection from intense UV radiation (Remias & Lutz, 2007) and helps to ensure liquid water availability in low-temperature environments (Dial et al., 2018), reinforcing a melt–albedo feedback in which higher algal abundance further accelerates snowmelt (Ganey et al., 2017).

Altitudinal zonation of biota in mountain regions, which was first documented by Humboldt and Bonpland in 1809, describes the layering of ecosystems that emerge as temperature, humidity, and solar radiation shift with elevation (Von Humboldt & Bonpland, 2010). A key transition is the tree line, which marks a gradual boundary between montane and alpine environments and limits the upper range for large plants to grow (Körner, 2012). Above this level, forested landscapes transition into subnival zones characterized by sparse vegetation, increased wind exposure, and progressively more persistent snow cover, ultimately giving way to the nival layer. This shift also seems to correspond to changes in the habitats where snow algae occur. Observations of *S. nivaloides* suggest that it mainly occurs above the tree line where snow persists for several months into the melt season. For example, across five years of satellite observations covering 94,200 km^2^ of the European Alps, red blooms dominated by *S. nivaloides* occurred predominantly above ∼1,800 m a.s.l., revealing their general absence from lower-elevation snowpacks (Roussel et al., 2024). Consistent with this high-elevation pattern, red blooms historically described as *Chlamydomonas nivalis* and now largely attributed to *S. nivaloides* have been reported from penitentes fields in the Andes at elevations above 5,200 m a.s.l. (Vimercati et al., 2019). By contrast, *Chloromonas* and *Chlainomonas* occupy a broader ecological niche including above tree line, below the tree line into shaded or low-light environments, or lake margins or ice-covered shorelines (Hoham, 1974; Procházková et al., 2018; 2019b; Engstrom et al., 2020). These observations suggest that elevation-associated habitat transitions may structure the distribution of specific snow algal taxa.

Elevation shapes the physical and radiative environment of mountain snowfields that likely impact snow algae (Frei & Schär, 1998; Takeuchi et al., 2013). For instance, UV radiation intensifies with elevation (Blumthaler et al., 1997), while colder air reduces snow water content (Rice et al., 2011). Snow density, metamorphic processes, and atmospheric deposition also vary by elevation. Shifts in dominant algal taxa across strong environmental gradients in glacier systems, often reflecting transitions between snow and ice communities, have been documented in the Himalayas, Alaska, and the Andes (Yoshimura & Kohshima, 1997; Takeuchi, 2013; Takeuchi & Kohshima, 2004). In these studies, algal biomass generally declined with elevation, where colder, drier conditions restrict meltwater availability and limit bloom formation at higher elevations (Yoshimura & Kohshima, 1997; Takeuchi, 2013; Tanaka et al., 2016). However, these patterns largely reflect shifts across the snow–ice boundary and differences in habitat preference between snow and ice algae, rather than elevational responses within snow algal communities. Data are needed on the impact of elevation on algal composition and traits— such as pigment investment, cell size, and biomass—to understand and quantify how these influence snow albedo. The steep increase in UV exposure with elevation further raises questions about whether algae accumulate more protective pigments, whether blooms intensify in color, and whether such adjustments amplify their contribution to albedo reduction across mountain snowfields.

Here, we hypothesized that elevation influences the abundance and optical properties of snow algae primarily through physiological and cellular adjustments, particularly shifts in pigment composition, and that these changes ultimately shape snow surface reflectance. Specifically, we predicted that with increasing elevation: (i) snow algae would exhibit increased investment in photoprotective pigments, particularly astaxanthin; (ii) cell size would shift toward larger cells under the increased environmental stress of higher elevation; and (iii) these physiological changes would intensify their contribution to snow albedo reduction. Our goal was to evaluate how these drivers interact along a broad altitudinal gradient using combined spectral, chemical, and biological measurements. Because our dataset spanned two latitudinal transects of mountain ranges (Cascade Range and Rocky Mountains), we considered two levels of analysis: the entire data set and each transect in isolation. This approach allowed us to assess whether the factors controlling algal effects on snow albedo are consistent across mountain ranges or shaped by regional context.

## Methods

### Snow sampling and processing

Sampling was designed to account for latitudinal differences in snowpack persistence (**Figure S1**). At lower latitudes (37–42° N), snow at high elevations tends to melt earlier than at higher latitudes (47–48° N). Accordingly, sampling at the beginning of each field season was focused on more southerly regions and progressed northward as the melt season advanced, a pattern that was more consistently followed in the Rocky Mountains. However, given the large geographic extent of the study, the sampling strategy remained flexible, and additional sites were included opportunistically when access allowed during transit between regions, particularly in the Cascades.

We sampled three blooming plots (spanning a range of pinkish hues) and one visually defined non-blooming plot (white snow, with no apparent algal presence) from ∼100 mountain snowfields across seven states in the northwestern U.S.A. during the summers of 2022–2024, with sampling conducted predominantly above the tree line or in open high-altitude areas (**Table S1**). Each plot was collected by removing the upper 7.5 cm of snow from a 36 × 36 cm quadrant using an avalanche shovel. The snow was placed in a clean bag, gently homogenized, and transferred to sterile 3.8-L Ziploc bags. Samples were kept cool and dark and transported directly to a nearby laboratory or field station, where they were processed within 3–6 h whenever possible.

Snow was melted gradually and passed through an acid-washed (10% HCl v/v) 125-μm Nitex nylon mesh to remove coarse particles. The filtrate was collected on pre-combusted (4.5 h at 500 °C) and acid-washed Whatman GF/C filters for analysis of particulate organic carbon (PC) and nitrogen (PN), and pigments, while MCE filters were used for DNA sequencing. Subsamples were also preserved with Lugol’s solution (6.3% v/v) and stored in the dark for subsequent microscopic counts and cell sizing.

### Particulate carbon (PC) and nitrogen (PN)

Particulate C and N were analysed by cutting each filter in half, with one half used for C and the other for N analysis. For C, filters were rinsed with 1 M HCl to remove carbonate minerals, then triple rinsed with 18.2 MΩ cm^−1^ deionized water and dried (8 h at 60 °C). Filters for N analysis were dried but not acidified. Subsequently, filters were placed into tin boats, sealed, and analyzed via a Costech Instruments Elemental Analyzer (EA) connected to a Thermo Scientific Delta V Advantage Isotope Ratio Mass Spectrometer (IR-MS) at either the Stable Isotope Facility (SIF) at UC Davis (U.S.A.) or the Metals Environmental and Terrestrial Analytical Laboratory (METAL) at Arizona State University (U.S.A.).

### Cell density and cell volume estimates

Cell density was determined using an inverted microscope (Nikon Eclipse E400 with 10X objective) equipped with a Sony camera (Alpha6000). Image software Fiji (ImageJ, version 1.54) was used to view images and manually count cells, following Schindelin et al. (2012). We estimated snow-algal cell volumes for ∼10 randomly selected samples per mountain range, chosen to best span the altitudinal gradient. The cell volume was obtained by measuring the major diameter of 18-24 cells from each sample using a light microscope (Leitz LaborLux S with 10x objective) to compute the cell volume assuming a spherically shaped cell. Analyses focused on *Sanguina nivaloides*-dominated samples, for which the spherical approximation is appropriate.

### DNA extraction

DNA was extracted from the MCE filters using a DNeasy PowerSoil Kit (Qiagen, Carlsbad, CA, USA) according to the manufacturer’s instructions and Almela and Hamilton (2025b). Negative controls were included: extraction blanks to check for kit contamination and field blanks. Field blanks consisted of MCE filters rinsed with ultrapure water (18.2 MΩ) and processed using the same equipment and procedures as the samples. The concentration of DNA was determined using a Qubit DNA Assay kit (Molecular Probes, Eugene, OR, USA) and a Qubit 3.0 Fluorometer (Life Technologies, Carlsbad, CA, USA). No DNA was detected in the negative controls.

### 18S rRNA amplicon sequencing and analyses

DNA was submitted to the University of Minnesota Genomics Center (UMGC) for sequencing using a Nextera XT workflow, 2×300bp chemistry, and the primers 1391f and EukBr that target the V9 region of 18S SSU rRNA (Amaral-Zettler et al., 2009; Stoeck et al., 2010).

Diversity and composition were assessed with QIIME v2-2024.10 (Bolyen et al., 2019). Briefly, cleaned and trimmed paired reads were filtered and denoised using the DADA2 plug-in (Callahan et al., 2016). For chimera identification, 340,000 training sequences were used. Identified operational taxonomic units (OTU), defined at 97% of similarity, were aligned using MAFFT (Katoh et al., 2002) and further processed to construct a phylogeny with fasttree2 (Price et al., 2010). Taxonomy was assigned to OTUs using the q2-feature-classifier (Bokulich et al., 2018) and blasted against the SILVA v138 99% 18S sequence database (Quast et al., 2012). All non-Chlorophyta sequences were removed from the dataset. Sequences assigned to Chlorophyta in SILVA were manually validated using BLASTN to confirm or refine taxonomic assignments. For each query, we retained the top-scoring match with 100% query coverage, requiring a minimum sequence identity of ≥99.4% to be considered a database match (consistent with Lutz et al., 2019). Sequences below this threshold were classified as “no BLAST hit.”. When the top BLAST hit corresponded to an “uncultured eukaryote” or “unknown Chlorophyta”, we selected the next best alternative hit based on the highest maximum score to improve the ecological interpretability of our dataset.

The 50 most abundant OTUs across all samples, including taxonomic assignments and BLAST statistics, are presented in **Table S2**. In addition, OTUs sharing the same taxonomic assignment were grouped to estimate the relative abundance of each algal taxon within individual samples; these community composition data are provided in **Table S3**.

### Pigment extraction and analysis

To characterize and quantify the major pigments in algal cells, a filter with algal biomass was placed in sterile 15-mL conical tubes, and 5 mL of a 7:2 acetone:methanol solvent was added. Cell disruption was performed by sonication (50% amplitude, 2 minutes). Samples were stored at -20 °C overnight and then sonicated again (50% amplitude, 1 minute) and centrifuged (5,000 × g, 4 °C, 10 min). An aliquot of the supernatant was filtered through a Millex® PVDF syringe filter (0.22 μm pore size). The filtered extracts were stored in glass vials at -80 °C until further analysis.

To quantify pigment concentrations, 10 μL of total extract was analyzed with an Agilent 1100 series HPLC using a Discovery® 250 × 4.6 mm C18 column (Sigma-Aldrich) with an isocratic mobile phase consisting of 50:20:30 methanol-acetonitrile-ethyl acetate flowing at 1 mL·min-1. Pigments were identified based on retention time and spectral matching to authentic standards of astaxanthin (3.17 min, 480 nm), chlorophyll-b (3.91 min, 649 nm), chlorophyll-a (4.39 min, 662 nm), and β-carotene (6.90 min, 454 nm). Diagnostic wavelengths for chlorophylls-a and -b were chosen to prevent interference with astaxanthin. The amount of compound in each peak was determined by integrating peak areas using OpenLab software (Agilent) and comparing values to those from standard curves prepared for each pigment.

Pigment measurements were calculated in volumetric units of melted snow. Per-cell pigment contents were then estimated by normalizing volumetric pigment concentrations by the total algal cell abundance in each sample.

### Vertical snow sampling for chlorophyll analysis

Vertical snow samples for chlorophyll analysis were collected using a 2.5-cm diameter corer, separating the snowpack into three consecutive layers (0–2.5 cm, 2.5–7.5 cm, and 7.5–12.5 cm). Samples were melted and filtered through pre-combusted Whatman GF/F filters, which were stored frozen (–20 °C) until analysis. Chlorophyll-a was quantified after overnight extraction in 90% acetone using the acid-correction method (EPA Method 445.0) on a Turner 10-AU fluorometer (Optical Kit #10-037R).

To assess the vertical distribution of biomass within the snowpack, we calculated the Upper 2.5-cm Chl-a fraction, defined as the proportion of total chlorophyll-a contained in the uppermost 2.5 cm layer relative to the integrated chlorophyll-a across the full 12.5 cm snow column.

### Measurements of snow physical properties and albedo

In the field, and adjacent to each measurement site, snowpack depth was recorded using an avalanche probe. Snow density was estimated from the 12.5-cm cores collected for vertical biomass profiling as follows: after melting each core, the meltwater volume was measured, and snow density was calculated as meltwater volume divided by the original core volume.

Prior to snow excavation, albedo was measured across wavelengths from 350 to 2,500 nm using an ASD FieldSpec® 4 hyperspectral spectroradiometer (Malvern Panalytical, USA) equipped with a cosine collector. This configuration integrates reflected radiation over a 180° field of view, providing hemispherical reflectance (i.e., albedo). The cosine receptor was mounted on a 100 cm pole and held level at ∼80 cm above the surface using a bubble level, oriented south to avoid casting a shadow on the target. At this height, the cosine receptor has an approximate field of view with a 35 cm radius, closely matching the sampled surface area. All measurements were conducted within ±3 h of solar noon under clear-sky conditions, and triplicate measurements were taken at each sampling point approximately 5 s apart. Measurements were acquired by standing down-sun of the sensor and recording sequential scans of downwelling and upwelling radiation. Because albedo was calculated as the ratio of consecutive measurements, no irradiance calibration was applied. Albedo measurements were not obtained for all samples and depended on instrument availability and suitable meteorological conditions during field sampling.

### Data analysis and statistics

Analyses of community composition and its variation with elevation were conducted using the full dataset (n = 294 samples), including both blooming (n = 206) and non-blooming (n = 88) conditions. Analyses focused on bloom-specific patterns were restricted to the subset of visibly red algal blooms (n = 206). For subsequent analyses of biomass, cellular traits, and optical properties, the dataset was further restricted to samples where S. nivaloides accounted for >55% of total algal sequences (n = 117), to isolate intraspecific physiological patterns from shifts in community composition. A low-elevation sample in which Sanguina was abundant (1,288 m a.s.l.) was excluded from this subset to avoid undue influence of a single low-elevation point, resulting in a reduced elevational range (1,876–3,423 m a.s.l.) for Sanguina-focused analyses.

All statistical analyses were conducted primarily in the R software package (v.4.x). To evaluate whether bloom developmental stage could act as a confounding factor, we used astaxanthin per cell and the astaxanthin:chlorophyll-a ratio as proxies of bloom maturity, based on the expectation that astaxanthin accumulates as Sanguina cells transition from green vegetative stages to red resting cysts. We then tested whether these proxies varied with sampling day-of-year (DOY) and snow depth, to represent the temporal progression of blooms. Differences in DOY and snow depth across regions and along the latitudinal gradient were assessed using linear models, whereas their relationships with astaxanthin per cell and the astaxanthin:chlorophyll-a ratio were evaluated using linear mixed-effects models fitted with the lme4 package in R, including site (snow patch) as a random intercept.

To explore general patterns of covariation among elevation and biological and environmental variables, pairwise relationships were evaluated using Pearson correlations with pairwise deletion of missing values. Before analysis, outliers were identified and removed using the ROUT test (Q = 0.2%) in GraphPad Prism (v.8.0.1.244). Outlier detection was performed separately for each region-specific subset to account for context-dependent variability, and the filtered data were then combined for analyses of the full dataset. For each comparison, we extracted the correlation coefficient (r) and associated P value. To control for multiple comparisons, Bonferroni corrections were applied separately to predefined groups of variables representing distinct processes: (i) biological variables (algal cell density (log_10_ cells mL^−1^), PC (µM mL^−1^), astaxanthin (µg L^−1^), chlorophyll-a (µg L^−1^), and pigment ratios), (ii) environmental variables (water content, estimated using the Normalized Difference Water Index (NDWI; Gao, 1996), snow density (g cm^−3^), PN (µM mL^−1^), and PC:PN ratios, and (iii) albedo metrics (bloom albedo, cell-normalized albedo, and relative albedo). Cell volume was excluded from this analysis because we had multiple replicates per sample.

To identify the main drivers of snow reflectance, we fitted linear mixed-effects models using spectral reflectance (full range: 350–2,500 nm) as the response variable. Site (snowfield) was included as a random intercept to account for the non-independence of samples collected within the same snowfield. Candidate predictors included environmental controls (latitude, longitude, elevation (m a.s.l.), snow depth (cm), snow density (g cm^−3^)) and biological controls (algal cell density (log_10_ cells mL^−1^), PC (µM mL^−1^), astaxanthin (µg L^−1^), chlorophyll-a (µg L^−1^), and the percentage of chlorophyll-a in the top 2.5 cm of snow). Water content (NDWI) was excluded because it is directly derived from the reflectance spectrum and would therefore introduce circularity. Cell volume was also excluded because it is an average of multiple measurements per sample and was available only for a subset of samples. Prior to model fitting, multicollinearity was assessed using variance inflation factors (VIF) from the car package, and variables with VIF > 3 were removed. Continuous predictors were standardized to improve model convergence and comparability of effect sizes. We constructed separate model sets to examine biological and environmental contributions to reflectance. Models were first fitted using *Sanguina*-dominated blooms to isolate physiological and snow-structural drivers within a single taxon, and subsequently extended to the full dataset, including non-*Sanguina* communities and snow without visible algae, to capture the full optical gradient from clean to biomass-rich snow. Because optical drivers may differ between mountain systems, region-specific models (Cascade Range and Rocky Mountains) were also constructed following the same variable-selection and collinearity criteria.

## Results & Discussion

### 1. Community composition

#### 1.1. Community composition across elevation and regions

We collected 294 samples in the Cascade Range and Rocky Mountains. Sample sites included broad and overlapping altitudinal range (1,059–3,423 m a.s.l.), enabling assessment of elevation-dependent patterns independent of geographic location as well as consideration of distinct mountain regions. Our study area also spanned a large latitudinal range from 37.7° to 48.9° N (**Table S1**). At our study sites, snow was sampled from visibly red algal blooms (n=206) and from adjacent visually defined white, non-blooming snow (n=88), allowing comparisons of community composition between bloom and non-bloom conditions.

Across all samples, *Sanguina nivaloides* was the most abundant algal taxon in 143 of 294 samples, regardless of elevation. On average, *S. nivaloides* represented 41.4 ± 39.2% of total algal sequences, followed by *Chlainomonas* sp. (12.7 ± 25.4%), *Chloromonas alpina* (5.3 ± 14.6%) and *Sanguina* sp. (0.7 ± 4.0%), while algal sequences with ‘no blast hit’ averaged 34.4 ± 34.9% (**Table S3**). Consistent with this pattern, *S. nivaloides* accounted for a substantial fraction of the most abundant OTUs (28 of the top 50; **Table S2**), indicating its dominance across both relative sequence abundance and OTU-level representation.

Among the 206 sequenced bloom samples (i.e., snow exhibiting visibly reddish algal blooms), *S. nivaloides* was the most abundant taxon in 128 samples (mean relative abundance of 51.9 ± 38.5%; **Figure 1**) followed by *Chlainomonas* sp. (15.1 ± 27.6%) and *C. alpina* (4.4 ± 12.7%). Together, these genera represent the principal taxa driving snow algal blooms (Hoham & Remias, 2020; Engstrom et al., 2020). Algal sequences with ‘no blast hit’ averaged 27.5 ± 27.7%.

**Figure 1.**
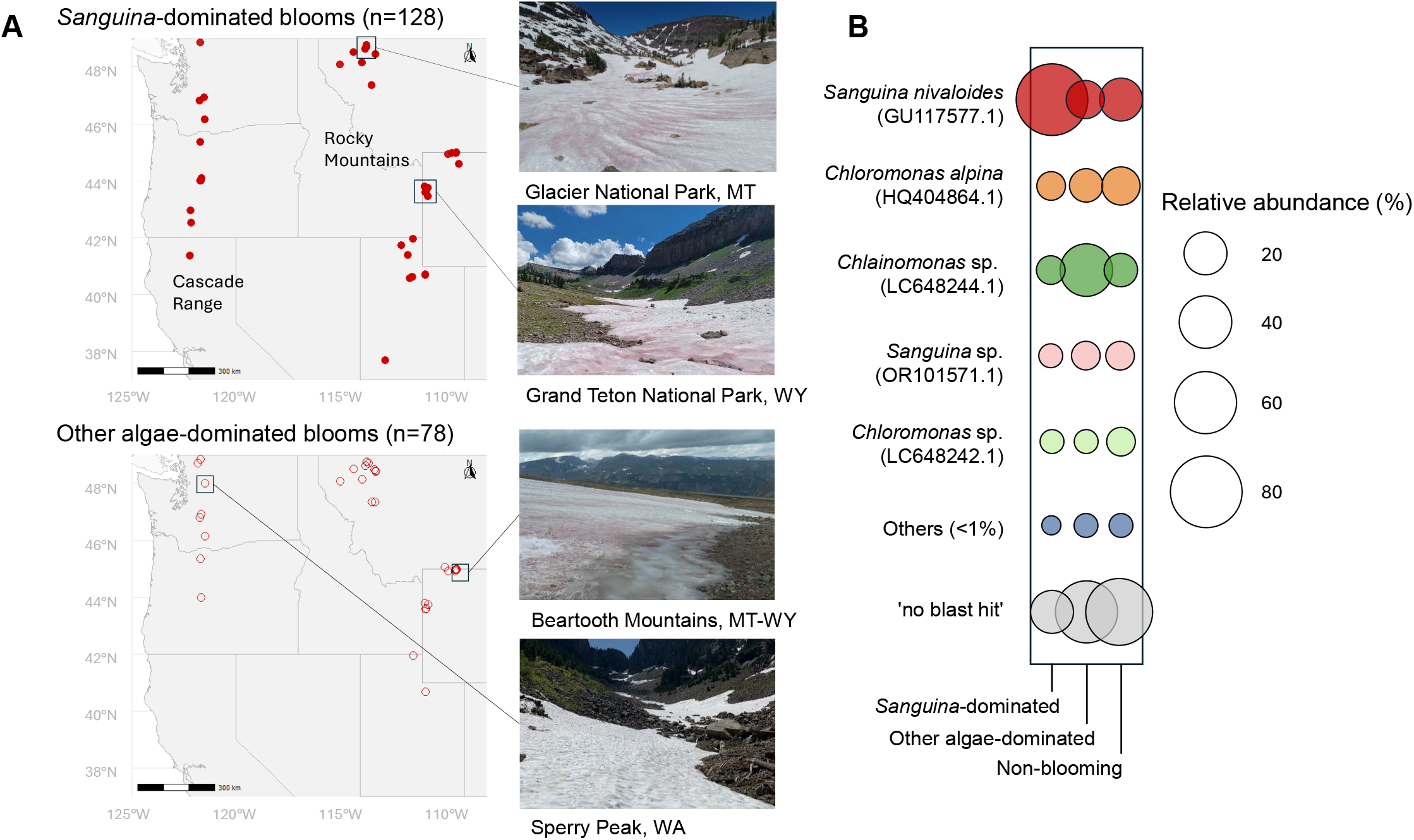
(**A**) Snow algae samples collected for this study during the summers of 2022–2024 across the Cascade Range and Rocky Mountains, separated by bloom type. The upper map shows *Sanguina*-dominated blooms (i.e., samples where *Sanguina nivaloides* was the most abundant taxon in visibly pigmented snow), whereas the lower map shows samples dominated by other snow-algae taxa. At each snowfield, an additional sample was collected from adjacent snow with no visible algal presence (“no visible algae”), which is not shown on the maps. Representative photographs from three sampling locations illustrate the range of sampled snowfields. (**B**) Algal community composition of samples classified as ‘*Sanguina*-dominated’, ‘Other algae-dominated’, and ‘Non-blooming’ (visually identified as white snow with no visible algal presence), based on 18S rRNA amplicon sequencing.

#### 1.2 Context-dependent dominance of snow algal taxa

Across the full elevational range sampled (1,059–3,423 m a.s.l.), the relative abundance of snow algal OTUs (**Table S3**) showed no significant association with elevation (*P* > *0*.*05*). This lack of a clear compositional shift contrasts with earlier reports of vertical zonation in snow algal communities along mountain gradients. Such zonation, often involving shifts in dominant taxa between ice- and snow-dominated habitats, has been reported from glaciers in the Himalayas (Yoshimura & Kohshima, 1997), Alaska (Takeuchi, 2013), and the Andes (Takeuchi & Kohshima, 2004). In these systems, taxonomic turnover has been attributed to strong local environmental gradients, including differences in snowpack stability, meltwater availability, light exposure, and nutrient supply. In contrast, our dataset includes mostly seasonal snowfields sampled during the melt season from ∼1,100 - 3,400 m a.s.l. Across this range, UV irradiance increases with elevation, whereas temperature and melt conditions in seasonal snow are more spatially heterogeneous and less systematically structured than in studies spanning transitions from seasonal or perennial snow to glacier ice. Under our sampling conditions, fine-scale zonation is less evident, and broader ecological patterns emerge, reflecting the capacity of dominant snow algal taxa to persist across a wide range of physical conditions in mountain snowfields above the tree line. Because our sampling was conducted predominantly above the tree line or in open high-altitude areas, we likely did not capture forested habitats where distinct snow algal communities may occur (Hoham, 1974; Procházková et al 2018; Procházková et al. 2019b; Engstrom et al., 2020), which may explain the absence of a clear transition in community composition along the elevational gradient in our dataset.

*S. nivaloides* was abundant in most of our samples above 1,800 m a.s.l. This pattern is consistent with previous observations from the European Alps, where red snow blooms dominated by *Sanguina* are preferentially associated with high-elevation snowpacks (Roussel et al., 2024). Similarly, on Gulkana Glacier in Alaska, red snow blooms historically attributed to *Chlamydomonas nivalis* were reported to dominate at higher elevations (Takeuchi, 2001); based on phylogenetic evidence, this organism is now generally recognized as corresponding to *S. nivaloides*, the primary organism associated with red snow (Procházková et al., 2019a).

Among the 206 sequenced bloom samples, at lower elevations (<1,500 m a.s.l.; n = 12), *Chlainomonas* sp. (27.3 ± 36.3%) and *C. alpina* (21.7 ± 36.3%) were more abundant. In our data set, all sub-1,500 m a.s.l. samples were collected from Mt. Baker–Snoqualmie National Forest, WA (**Figure 1**). Previous studies have reported seasonal blooms of *Chlainomonas* in snow-on-lake sites, with some members of this genus preferentially occurring in wet snow (e.g., Novis et al., 2008; Procházková et al., 2018; Beck, 2024; Matsumoto et al., 2024). The higher relative abundance of *Chloromonas* and *Chlainomonas* in our lower-elevations sites at Mt Baker could reflect association with lake environments and wetter snow conditions, although proximity to forested areas below the tree line may also contribute. Both genera also dominated a limited number of individual samples at elevations above 2,000 m a.s.l., and in some cases above 3,000 m a.s.l.; however, this dominance was not consistent across replicate samples from the same bloom, indicating substantial within-bloom heterogeneity in community composition In contrast, *S. nivaloides* was consistently dominant across most samples, particularly at higher elevations, indicating contrasting patterns between lower- and higher-elevation sites. However our low-elevation dataset is geographically restricted and small, limiting our ability to assess the role of elevation relative to local habitat-specific conditions in structuring snow algal community composition.

### 2. Variation in *Sanguina* abundance and biomass distribution

#### 2.1. Altitudinal variation in biomass and cellular traits

*S. nivaloides* was abundant in most of our samples that exhibited visibly reddish blooms. We therefore subset the data to sites where *S. nivaloides* accounted for >55% of total algal sequences (n = 117), with a mean relative abundance of 82.6 ± 12.0% within this subset. This approach allowed us to examine altitude-related trends in biomass, cell size, and pigment composition as indicators of intraspecific physiological acclimation in *S. nivaloides*, rather than shifts driven by community turnover (**Figure 1A**).

For this subset of samples, snow algal cell density varied widely (mean = 17,938 cells mL^−1^; range: 1,667–147,500 cells mL^−1^; **Table S1**), but showed no significant relationship with elevation (**Figure 2**), contrasting with observations from Himalayan and Arctic glaciers, where cell density declined at high altitude (Yoshimura & Kohshima, 1997; Takeuchi, 2013). While lower-elevation snowpacks may provide wetter conditions that favor rapid algal growth, they are typically short-lived. In contrast, snow at higher elevations persists for longer periods, potentially allowing sustained algal accumulation despite colder temperatures and shorter daily growth windows. These opposing effects may result in no clear elevational trend in cell density.

**Figure 2.**
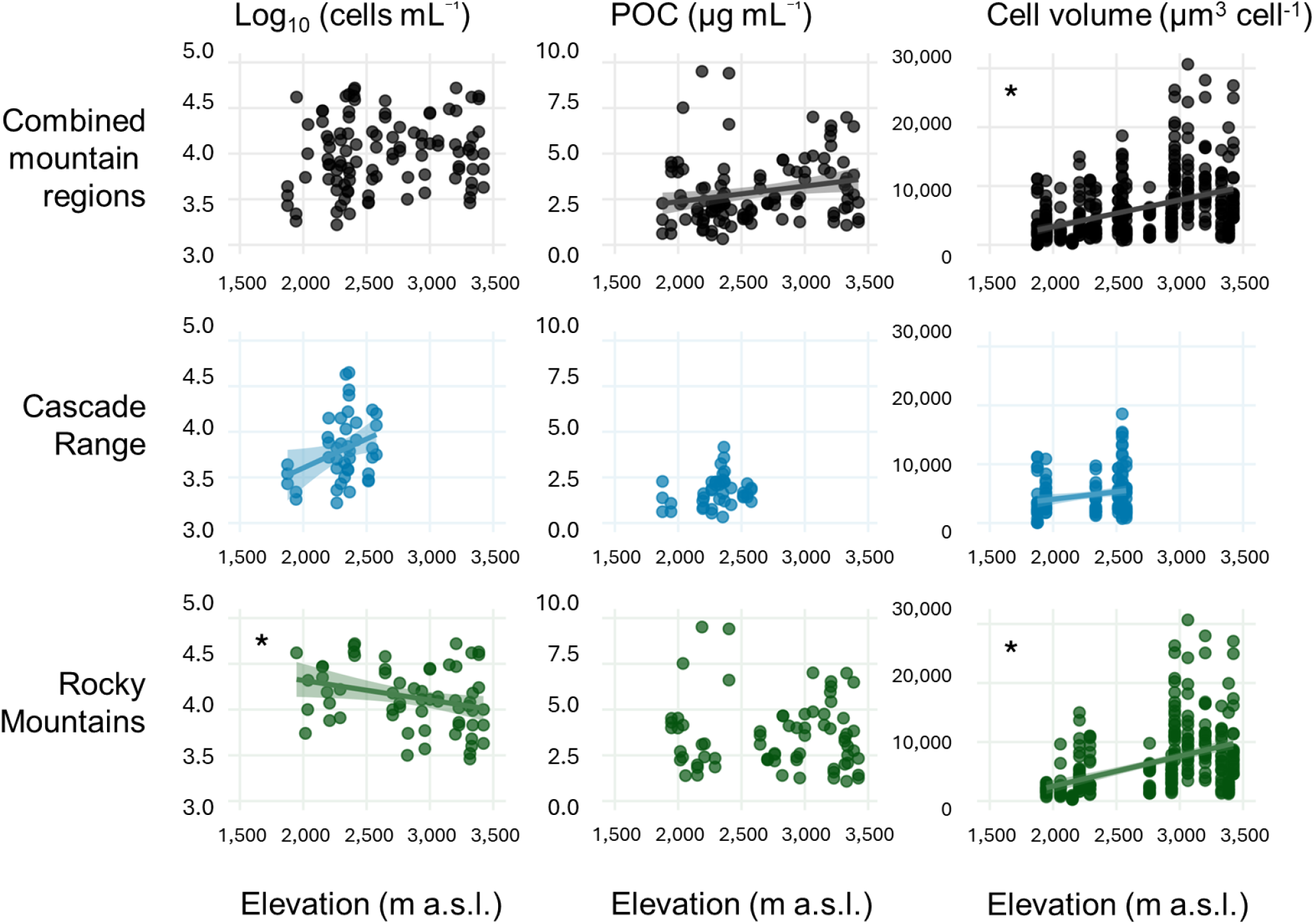
Relationships between elevation and biotic variables for the combined dataset and each mountain range (rows: Combined, Cascade Range, Rocky Mountain Range). Variables include cell density (cells mL^−1^), particulate organic carbon (PC, µg mL^−1^) and cell volume (µm^3^ cell^-1^). Points represent individual observations after outlier removal; solid lines indicate linear relationships significant at *P* < *0*.*05* (95% confidence intervals shown). Asterisks (*) denote relationships that are significant after Bonferroni correction (*P* < *0*.*01*).

**Figure 3.**
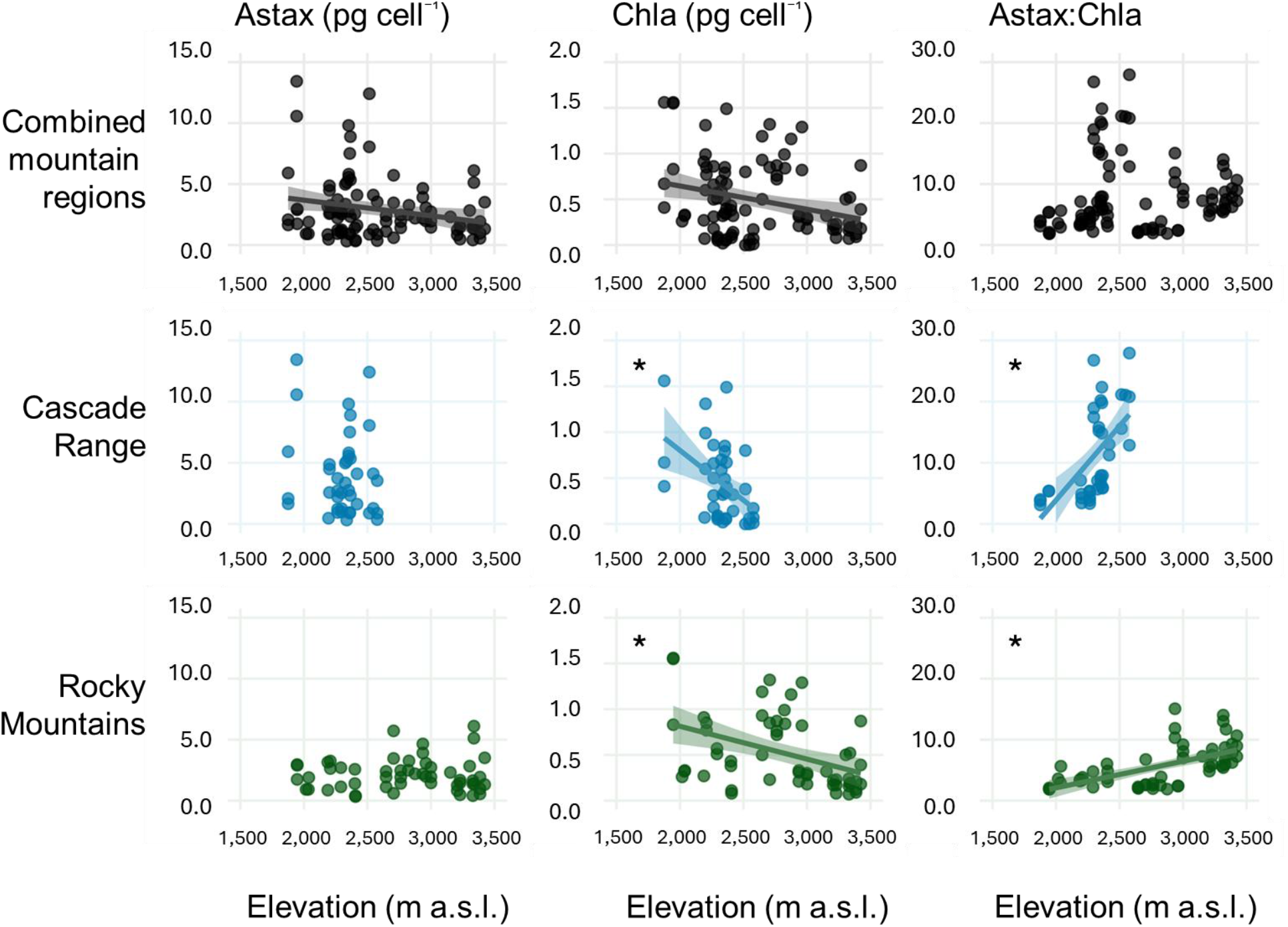
Relationships between elevation and biotic variables for the combined dataset and each mountain range (rows: Combined, Cascade Range, Rocky Mountain Range). Variables include astaxanthin concentration (Astax, µg L^−1^), chlorophyll-a concentration (Chla, µg L^−1^), astaxanthin per cell (Astax, pg cell^−1^), chlorophyll-a per cell (Chla, pg cell^−1^). Points represent individual observations after outlier removal; solid lines indicate linear relationships significant at *P* < *0*.*05* (95% confidence intervals shown). Asterisks (*) denote relationships that remain significant after Bonferroni correction (*P* < *0*.*01*).

Instead, mean cell volume increased with elevation from 3.30 ± 2.75 µm^3^ per cell below 2,000 m a.s.l. to 8.92 ± 6.22 µm^3^ per cell above 3,000 m a.s.l. (r = 0.44; *P* < *0*.*01*). Particulate organic carbon (PC) concentrations in snow algal blooms also increased with elevation from 2.73 ± 1.69 µM C mL^−1^ to 3.76 ± 1.82 µM C mL^−1^ (r = 0.22; *P* < *0*.*05*), suggesting a weak positive trend that was not significant after Bonferroni correction (**Figure 2; Table S4**). Together, increasing cell size and PC indicate greater biomass per cell at higher elevations.

Along the altitudinal gradient, PC:PN ratios showed no consistent trend, remaining within a relatively narrow range (mean = 23.51 ± 10.31; 20.08 ± 15.76 below 2,000 m a.s.l. and 22.79 ± 8.34 above 3,000 m a.s.l.). This stability indicates that bulk stoichiometry remains largely unchanged with elevation and does not reveal a clear shift in nutrient status along the gradient at the bloom scale, consistent with previous enrichment experiments showing no biomass response to nutrient additions (Hamilton and Havig, 2023; Millar et al., 2024; Klip et al., 2026).

We interpret increasing cell size as a response to environmental stress with elevation, including stronger UV radiation and lower air temperatures. Similar morphological plasticity has been reported in other snow algae, such as Raphidonema nivale, which alters cell size and shape in response to environmental conditions (Stibal & Elster, 2005). Although nutrient stress can also promote increases in cell size (Almela et al., 2024), our results do not support a dominant role of nutrient limitation in structuring these patterns.

Given that cell abundance showed no consistent relationship with elevation, we considered whether alternative strategies might mitigate high-altitude stress, including vertical positioning within the snowpack (Painter et al., 2001; Ono & Takeuchi, 2025; Ono et al., 2025). Snow algae can reduce high-radiation exposure by occupying subsurface layers while still influencing surface albedo (Almela et al., 2025c). However, the Upper 2.5-cm Chl-a fraction showed no relationship with elevation, indicating no consistent elevational trend in biomass distribution within the upper 12.5 cm of the snowpack. In 45% of samples, the highest chlorophyll-a concentration occurred in the uppermost 2.5 cm layer (**Table S1**). Any elevational differences in vertical stratification, if present, therefore likely occur at finer vertical scales (sub-cm) at the snow surface that were not resolved by our sampling.

#### 2.2. Elevational shifts in pigment allocation

To assess how pigment composition varied with elevation, we quantified pigment per-cell pigment concentrations. Per-cell astaxanthin and chlorophyll-a contents declined with elevation (r = −0.22, *P* < *0*.*05* and r = −0.28, *P* < *0*.*01*, respectively), but neither relationship remained significant after Bonferroni correction (**Table S4**). The decline was more pronounced for chlorophyll-a, decreasing from 1.10 ± 0.52 pg cell^−1^ below 2,000 m a.s.l. to 0.26 ± 0.18 pg cell^−1^ above 3,000 m a.s.l., compared to astaxanthin, which decreased from 5.15 ± 4.50 to 1.97 ± 1.45 pg cell^−1^ over the same elevation range. Because mean cell volume increases with elevation, these declines are consistent with larger cells containing less pigment at higher elevations. When normalized to cell volume, astaxanthin and chlorophyll-a concentrations were ∼12-fold and ∼20-fold lower, respectively, above 3,000 m a.s.l. than below 2,000 m a.s.l. (**Table S5**), although this assessment is based on a subset of samples for which we had both pigment and cell volume data (n = 16). Consistent with these patterns, the astaxanthin:chlorophyll-a ratio increased with elevation, rising from 3.44 ± 1.45 below 2,000 m a.s.l. to 7.80 ± 1.80 above 3,000 m a.s.l., although this trend was not significant.

The pigment ratios observed here span a wide range that overlaps with values reported for snow algae across contrasting cryospheric environments. For example, astaxanthin:chlorophyll-a ratios in red blooms vary widely, from ∼3 in samples from Mt. Tateyama in Japan (Nakashima et al., 2021) to ∼20.9 in the Austrian Alps (Remias et al., 2005), ∼34 in Svalbard (Müller et al., 1998), and up to 56 (Halbach et al., 2025). This variability underscores the high variability in pigment investment among regions and snow settings (e.g., Halbach et al., 2022; Davey et al., 2019; Remias & Lutz, 2007). Pigment composition in snow algae is strongly structured by taxonomic identity and life stage (e.g., Nakashima et al., 2021; Almela et al., 2025a). We used astaxanthin content per cell and the astaxanthin:chlorophyll-a ratio to approximate the bloom developmental stage. Neither change systematically with time of sample collection (DOY) or snow depth (**Table S6**), suggesting that the trends reported here are not driven by differences in bloom maturity across samples. Elevational shifts in cell size and pigment allocation observed across *Sanguina*-dominated samples in our dataset suggest that altitude-related environmental gradients drive consistent changes in pigment allocation, even in the absence of significant trends in pigment concentrations. Snow algae are known to exhibit physiological plasticity including adjustments of their photosynthetic and photoprotective machinery in response to changes in irradiance, temperature, and snowpack properties (Procházková et al., 2019b; Müller et al., 1998). In our data set, the increase in astaxanthin relative to chlorophyll-a is primarily driven by the decline in chlorophyll-a with elevation. Together with the less pigment in larger cells, these patterns are consistent with physiological acclimation to intensified radiation and other stressors characteristic of high-altitude snowfields.

#### 2.3. Regional patterns across mountain systems

To evaluate whether the overall patterns we observed were consistent across mountain regions, we repeated the analyses for the Cascade Range (samples between 1,876 and 2,577 m a.s.l.) and Rocky Mountains (samples between 1,947 and 3,423 m a.s.l.) separately. In the Cascade Range, cell density showed a weak positive relationship with elevation that was not significant after Bonferroni correction (r = 0.33; *P* < *0*.*05*), whereas in the Rocky Mountains cell density decreased significantly with elevation (r = −0.42; *P* < *0*.*01*) and remained significant after correction (**Figure 2; Table S4**). Mean cell densities at the highest elevations did not differ significantly between regions (Cascades >2,300 m a.s.l., n = 25: 11,872 cells mL^−1^; Rockies >3,000 m a.s.l., n = 29: 16,656 cells mL^−1^; *P* > *0*.*05*). This result indicates that algal abundance depends on environmental conditions, likely reflecting how melt schedules, chemical conditions, and other snowpack properties vary between mountain systems, as well as variability at the scale of individual snow patches. This interpretation is consistent with observations from other sites where biomass peaks at mid elevations rather than showing a consistent increase or decline (Takeuchi et al., 2001).

Mean cell volume increased with elevation in both mountain ranges (Cascade: r = 0.20, *P* < *0*.*05*; Rocky: r = 0.46, *P* < *0*.*01*), although this relationship was not significant after Bonferroni correction in the Cascade Range. Chlorophyll-a per cell declined with elevation (Cascade: r = −0.45; Rocky: r = −0.42; *P* < *0*.*01*), while the astaxanthin:chlorophyll-a ratio increased (Cascade: r = 0.61; Rocky: r = 0.61; *P* < *0*.*01*; **Table S4**). Collectively, these results indicate that, while algal biomass may vary regionally and be influenced by environmental conditions, physiological responses to increasing elevation are consistent across mountain systems. These responses include larger cell size and a shift toward higher astaxanthin:chlorophyll-a ratios, reflecting changes in pigment balance driven by reduced chlorophyll-a per cell and consistent with physiological adjustment to environmental stress at higher elevations (Remias et al., 2005). PC:PN showed a weak but significant increase with elevation in the Rocky Mountains (r = 0.11; *P* < *0*.*01*; **Table S4**), but this pattern was not observed in the Cascade Range, indicating only minor, region-specific variation in biomass composition.

To further examine regional differences, we compared physiological responses across samples from the same elevation range (1,876 and 2,577 m a.s.l). In the Cascade Range, astaxanthin per cell declined with elevation (from 5.69 to 3.70 pg cell^−1^), while chlorophyll-a per cell decreased more sharply (from 1.20 to 0.20 pg cell^−1^), resulting in a pronounced increase in the astaxanthin:chlorophyll-a ratio. In contrast, in the Rockies, both astaxanthin (2.53 to 1.45 pg cell^−1^) and chlorophyll-a (1.31 to 0.52 pg cell^−1^) declined with elevation, leading to a more moderate shift in their ratio. Notably, these patterns in the Cascades are associated with a latitudinal gradient across elevation (46.8°N below 2,000 m a.s.l vs 41.4-42.5°N above 2,400 m a.s.l), whereas Rocky Mountain samples span a broader latitudinal range at both low and high elevations (41.4-48.6°N). This co-variation of elevation and latitude in the Cascade Range likely amplifies environmental gradients, as higher-elevation sites also experience greater potential solar irradiance due to their lower latitude, with mean summer all-sky shortwave irradiance increasing from 5.73 kWh m^−2^ day^−1^ at ∼47°N to 7.06 kWh m^−2^ day^−1^ at ∼42°N (∼23% increase; NASA POWER, 2001–2020), potentially reinforcing changes in radiation regime. In contrast, the broader latitudinal spread in the Rockies decouples these factors, providing a more isolated test of elevation.

### 3. Regional context for elevational variability in snowpack conditions

Snowpack physical properties, including liquid water availability, snow structure, and the deposition of abiotic light-absorbing particles (LAPs), influence both snow algal growth and their optical effects, as snow albedo reflects the combined influence of biotic and abiotic impurities within the snowpack (Skiles et al., 2018). Because these factors co-vary with elevation, we examined how snowpack properties change with elevation across regions.

In our study, mean snow water content, estimated from the Normalized Difference Water Index (NDWI), was higher in the Cascade Range than in the Rockies (0.59 vs 0.53; t = 5.82, *P* < *0*.*01*; **Table S1**). A similar regional contrast was observed for snow density, which was higher in the Cascade Range than in the Rockies (0.58 vs 0.52 g cm^−3^; t = 3.91; *P* < *0*.*01*), consistent with wetter snow conditions in maritime regimes such as the Cascades (Trujillo and Molotch, 2014). Across the full dataset, NDWI showed no overall trend with elevation (**Table S4**). However, when analyzed by region, NDWI increased with elevation in the Cascade Range (r = 0.39; *P* < *0*.*01*) and showed a similar but non-significant tendency in the Rockies, indicating progressively drier snow at higher elevations. This pattern is consistent with the strong control of air temperature on snow liquid water content, whereby colder conditions limit melt–refreeze cycles and reduce water availability (Takeuchi, 2001; 2013; Yoshimura & Kohshima, 1997). In contrast, snow density and depth showed no consistent relationships with elevation, likely reflecting the integration of multiple local-scale processes—including precipitation regime, wind redistribution, avalanching, and topographic heterogeneity—that can mask simple altitudinal controls (Frei & Schär, 1998; Gauer, 2001; Lehning et al., 2008; Grünewald et al., 2014).

Abiotic LAPs may further modulate albedo–elevation relationships in seasonal snowfields. LAP deposition often decreases with elevation (Thomas & Duval, 1995; Takeuchi et al., 2013; Wittmann et al., 2017; Kavan et al., 2020) but localized mineral inputs from exposed soils and episodic dust events, such as those affecting high-elevation sites in the Rocky Mountains (Lang et al., 2023; 2025), may obscure simple elevational patterns. Because biotic and abiotic LAPs frequently co-occur and interact (Di Mauro et al., 2024), disentangling their individual contributions requires direct measurements beyond the scope of this study.

### 4. Elevation modulates the contribution of algae to snow albedo reduction

#### 4.1. Elevational patterns in algal-driven albedo

To assess how snow algae reduce albedo along the altitudinal gradient, we analyzed spectral reflectance (350–2,500 nm) in *Sanguina*-dominated blooms (hereafter, bloom albedo). When all samples were combined, bloom albedo showed no trend with increasing elevation, regardless of whether absolute values, cell density–normalized values, or relative albedo (bloom albedo vs. adjacent clean snow) were considered (**Figure 4A–C; Table S4**). In addition, normalizing albedo by cell volume did not yield a significant relationship with elevation, indicating that variation in cell size alone does not account for the optical patterns we observed.

**Figure 4.**
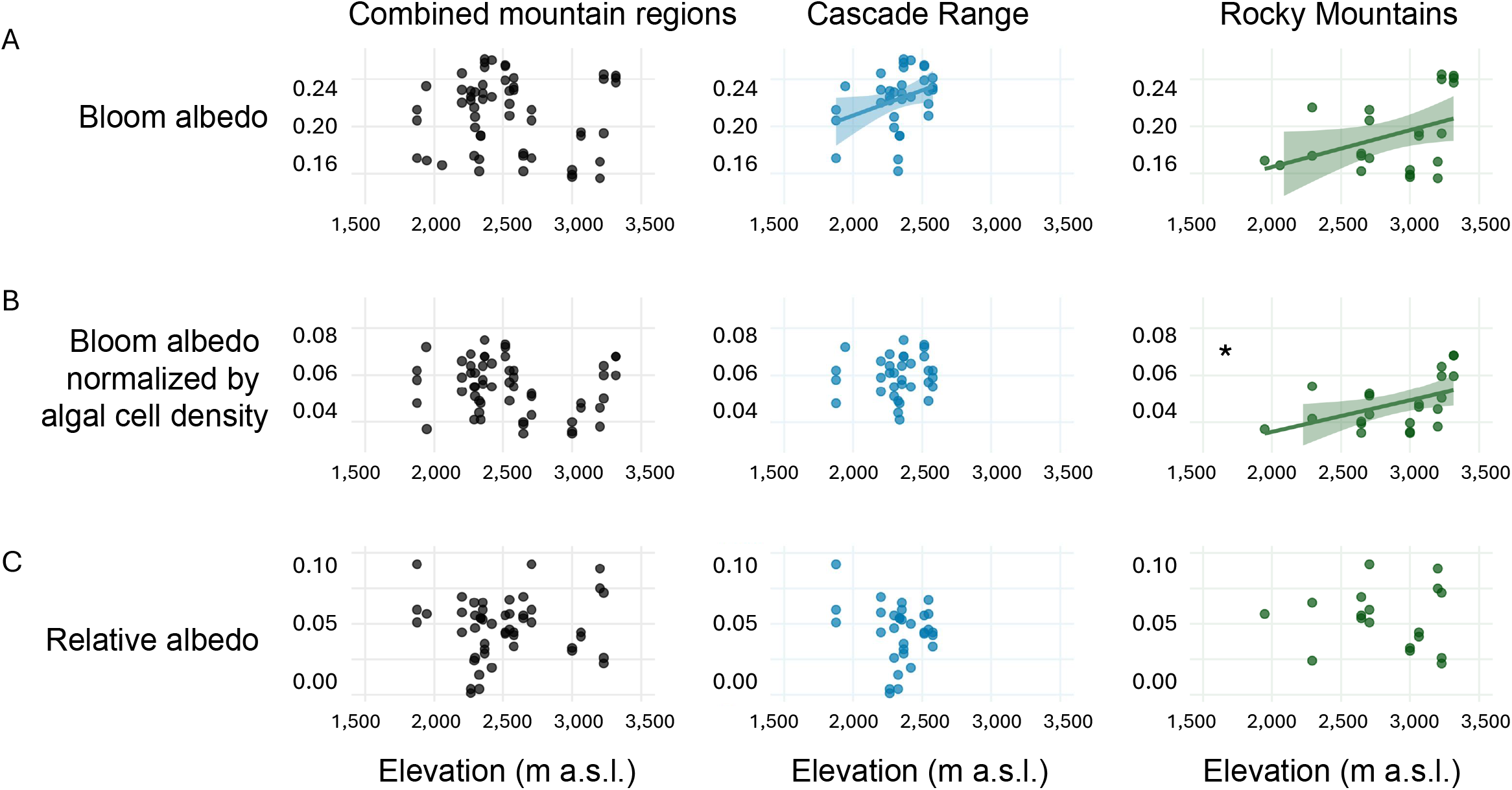
Elevational variation in snow optical properties across mountain regions. Snow albedo (350–2,500 nm) plotted as a function of elevation, for the combined dataset and for each mountain range separately (columns: Combined, Cascade Range, Rocky Mountains): (**A**) albedo in *Sanguina*-dominated bloom samples (i.e., bloom albedo), (**B**) bloom albedo normalized by cell density, and (**C**) relative albedo (bloom albedo vs adjacent clean snow albedo). In all panels, the x-axis shows elevation (m a.s.l.). Points represent individual observations after outlier removal; solid lines indicate linear relationships significant at *P* < *0*.*05* (95% confidence intervals shown). Asterisks (*) denote relationships that remain significant after Bonferroni correction (*P* < *0*.*01*).

To account for regional heterogeneity, we next analyzed the Cascade Range and Rocky Mountains separately. In contrast to the full data set, bloom albedo increased with elevation in both mountain regions, with statistically significant relationships in the Cascade Range and the Rocky Mountains (r = 0.35 and 0.41, respectively; *P* < *0*.*05*; **Figure 4A**), but these relationships were not significant after Bonferroni correction. Normalizing bloom albedo by algal cell density revealed a significant increase with elevation in the Rockies (r = 0.48; *P = 0*.*01*; **Figure 4B**), whereas no clear pattern was observed in the Cascades.

The contrasting results obtained when analyzing all samples together versus by region likely reflect the confounding influence of local snowpack heterogeneity on broader optical patterns and the inherent challenges of accurately measuring mountain snow albedo (Bair et al., 2018). An increase in absolute reflectance with elevation may result from drier snow conditions at higher elevations, increasing albedo regardless of the presence of snow algae. However, when bloom albedo was expressed relative to adjacent non-blooming snow (**Figure 4C**), its relationship with elevation differed from that observed for bloom albedo. In both mountain ranges, relative albedo values tend to be lower at higher elevations, whereas bloom albedo increased, although this contrast did not constitute a statistically significant elevational trend (**Table S3**). While tentative, this pattern is consistent with the possibility that accounting for background snow properties may reveal proportionally greater light absorption by high-elevation algal blooms, potentially reflecting enhanced pigment efficiency or packaging effects under harsher surface conditions at higher elevations.

#### 4.2 Physiological interpretation of elevational responses

Environmental constraints that vary with elevation shape algal physiology, thereby modulating their contribution to albedo reduction (**Figure 5**). At higher elevations, algal cells are exposed to increasing radiative stress, including stronger UV irradiance. Under clear skies, total solar irradiance increases by approximately 8 ± 2% per 1,000 m a.s.l., while UV radiation may rise at the studied latitudes by up to ∼18% per 1,000 m a.s.l. (Blumthaler et al., 1997). These conditions are expected to enhance oxidative stress through increased production of reactive oxygen species (ROS), which can damage cellular components and impair physiological function (Rezayian et al., 2019).

Astaxanthin:chlorophyll-a ratio increased with elevation in both mountain regions, indicating a relative shift toward photoprotective investment. Carotenoids can both mitigate and be susceptible to oxidative stress, acting as photoprotective antioxidants while also undergoing degradation under high ROS conditions (Sager & Zalokar, 1958; Bidigare et al., 1993). This process may contribute to the observed decline in per-cell astaxanthin with elevation, while the increase in the astaxanthin:chlorophyll-a ratio is primarily driven by a reduction in chlorophyll-a, suggesting a downregulation of photosynthetic investment under stress (Zheng et al., 2020).

Higher elevations are also characterized by lower air temperatures and repeated freeze–thaw cycles, which generally reduce snow liquid-water content (Rice et al., 2011), a key factor controlling algal bloom development. Snow algae typically proliferate when above-freezing conditions persist for several days, allowing meltwater to accumulate and persist (Hoham, 1980; Takeuchi, 2001; Roussel et al., 2024). Thus, snow algae at higher elevations are likely subject to a dual constraint of increased radiative stress and reduced liquid-water availability, consistent with previous observations of enhanced melt at lower-elevation sites (Yoshimura & Kohshima, 1997).

Under these combined stresses, our results indicate that populations of *S. nivaloides* respond through coordinated increases in cell size and shifts in pigment balance, resulting in higher astaxanthin:chlorophyll-a ratios despite lower pigment concentrations per cell. The decline in per-cell astaxanthin with elevation may therefore reflect increased pigment turnover or degradation under enhanced oxidative conditions. Within the Rocky Mountains, mean astaxanthin per cell increases from Montana to more continental sites in Wyoming and Utah (**Table S1**), consistent with greater solar exposure, although variability among sites is high, suggesting that these relationships may be context-dependent. While our dataset does not directly resolve pigment turnover, future work quantifying carotenoid degradation products could help clarify the role of oxidative stress in shaping pigment dynamics under high-elevation and high-UV conditions. These responses are consistent with the physiological plasticity documented in snow algae, whereby cells adjust morphology and pigment composition in response to environmental stress (e.g., Müller et al., 1998; Leya et al., 2009; Almela et al., 2024).

### 5. Integrative predictors of algal reflectance

#### 5.1 Drivers of albedo within *Sanguina*-dominated blooms

To disentangle the processes underlying variability in snow albedo, we modelled reflectance across the full spectral range (350–2,500 nm) using linear mixed-effects models that account for the non-independence of samples within snowfields. We first used the combined dataset to identify broad-scale controls on reflectance across the full environmental range captured in this study, and then performed region-specific analyses to assess the consistency of these patterns within mountain systems.

Across all *Sanguina*-dominated samples (**Table S7**), models including biological predictors showed a positive association between reflectance and astaxanthin concentration (t = 1.81), while other variables, including cell density and chlorophyll-a, exhibited weaker relationships (|t| < 1.7; n = 44). These effects remained modest after accounting for strong local variability, with site-level random effects explaining the majority of the variance (ICC ≈ 0.74). In contrast, models including environmental predictors revealed significant structure along the longitudinal gradient (t = −2.62) and a weaker association with elevation (t = 1.92), while snow depth and density showed no clear effects. As in the biological models, site-level variability remained dominant (ICC ≈ 0.78), indicating that reflectance is primarily structured at the scale of individual snowfields, with broader spatial gradients exerting a secondary influence.

Within individual mountain systems, contrasting patterns emerged. In the Cascade Range, reflectance showed no clear relationship with either environmental or biological predictors (|t| < 0.7 and < 1.5, respectively; n = 32), and was instead primarily structured at the snowfield level, with site-level random effects accounting for most of the variance (ICC ≈ 0.80–0.90), indicating strong local heterogeneity. In contrast, in the Rocky Mountains, reflectance showed clearer relationships with both biological and environmental predictors, decreasing with increasing cell density (t = −3.85) and increasing with astaxanthin concentration (t = 2.14), although the latter likely reflects shared responses to environmental conditions rather than a direct optical effect of the pigment. Models including environmental variables indicated positive associations with snow density (t = 3.64) and snow depth (t = 2.13). However, variance associated with site-level random effects could not be resolved in these models (singular fit; n = 12), consistent with limited power to detect between-snowfield variability. Despite this limitation, consistent relationships with both biological and snowpack-related predictors were observed, suggesting that reflectance may be more directly linked to biomass and snowpack properties in this region. Together, these results highlight strong regional differences in the controls on reflectance, with local heterogeneity dominating in the Cascade Range and more direct links to biological and snowpack properties emerging in the Rocky Mountains.

#### 5.2 Algal contributions to albedo across clean and bloom-covered snow

To evaluate algal effects on snow albedo beyond *Sanguina*-dominated blooms, we expanded our analysis to include other algae-dominated blooms and snow without visible algae (non-blooming samples), thereby capturing the transition from clean snow to biomass-rich surfaces (**Table S8**). Reflectance was primarily associated with total cell density (t = −2.16), while pigment-related variables showed no clear effects (|t| < 0.5; n = 54). Site-level variability remained substantial (ICC ≈ 0.71), indicating that local controls continue to dominate reflectance patterns. These results support algal biomass as a primary driver of biological albedo reduction in mountain snowpacks, consistent with previous observations across alpine and polar environments (Hotaling et al., 2021). They further indicate that reflectance is largely driven by total biomass rather than pigment composition, suggesting that differences in community composition or cell-level pigment adaptations do not override the dominant influence of biomass on snow darkening.

Models including snowpack and environmental predictors revealed consistent structure along the longitudinal gradient (t = −2.21) and a weaker association with elevation (t = 1.87), while snow depth and density showed no clear effects (|t| < 1.3; n = 54). As in the biological models, site-level variability remained substantial (ICC ≈ 0.67). Longitude emerged as a significant spatial predictor, likely capturing broad-scale contrasts in snowpack properties between the Cascade Range and the Rocky Mountains. Although our data document regional differences in snow water content and density, longitude likely integrates additional, unmeasured characteristics of maritime versus continental snow regimes, including temperature variability and accumulation dynamics (Trujillo and Molotch, 2014). This geographic signal indicates that algal impacts on albedo are modulated by regional snowpack context, potentially mediated by differences in snow physical structure or the co-occurrence of mineral LAPs. Such interactions may enhance albedo reduction beyond the effect of biological or abiotic components alone, as dust-enriched snow can both accelerate melt and stimulate algal growth (Hoham & Ling, 2000; Lutz et al., 2015), and combined effects of mineral particles and algal biomass can amplify darkening (Di Mauro et al., 2024).

Regional models revealed contrasting patterns between mountain systems. In the Cascade Range, reflectance showed no clear relationship with snow properties (|t| < 1; n = 41) and remained strongly structured at the snowfield level (ICC ≈ 0.64). Biological models revealed a clear negative association with astaxanthin concentration (t = −2.76), while other variables showed weaker effects (|t| < 1.5), and site-level variability remained high (ICC ≈ 0.79). In contrast, in the Rocky Mountains, reflectance showed clearer relationships with both physical and biological predictors. When assessing the snow properties in the models, results indicated a positive association with snow density (t = 2.66), while elevation showed a weaker trend (t = 1.57), with moderate site-level variability (ICC ≈ 0.66). The observed elevation–albedo relationship likely reflects covariation with other snowpack characteristics that change with altitude, such as grain size or liquid water content, as well as broader environmental gradients not explicitly captured in the model.

Biological models revealed strong relationships with both cell density and astaxanthin concentration, with reflectance decreasing with increasing cell density (t = −3.74) and increasing with astaxanthin (t = 3.05), although variance associated with site-level random effects could not be resolved (singular fit), indicating limited power to detect between-snowfield variability. The positive association with astaxanthin likely reflects shared responses to environmental conditions rather than a direct optical effect of the pigment itself. Notably, algal cell density showed no consistent relationship with elevation across regions, whereas cell volume increased with elevation, indicating that altitude structures cellular traits more strongly than algal abundance. Although cell size could not be explicitly incorporated into the models, this suggests that trait-mediated changes in light absorption, together with pigment investment, may contribute to snow darkening beyond biomass effects alone. Similar increases in cell size have been reported in other snow algal lineages (Novis et al., 2008), where larger cells tend to exhibit greater pigment accumulation and may reflect both physiological adjustment and longer-term adaptation to snow environments. Larger cells may also enhance light absorption per cell, strengthening the link between cellular traits and radiative effects.

Such mechanisms can enhance energy absorption and promote localized melting around cells, reinforcing feedback between pigmentation water availability (Ganey et al., 2017). In particular, secondary carotenoids such as astaxanthin are highly effective at absorbing visible radiation and dissipating excess energy as heat, promoting melt while limiting cellular overheating relative to other pigment colors (Dial et al., 2018). Because this confers an advantage under the high-UV, cold, low–liquid-water conditions characteristic of high-elevation snowfields, it may help explain the predominance of red blooms in high-alpine environments (Thomas, 1972; Almela et al., 2025a). Carotenoid production can also be induced by other stress factors, including nutrient limitation (Bidigare et al., 1993; Leya et al., 2009). However, the predominance of red over other pigment phenotypes in response to these stressors remains unresolved.

### 6. Broader hydrological relevance

In the western United States, seasonal snowpacks provide a major fraction of annual runoff and sustain streamflow through the dry summer months, contributing up to ∼70% in some mountain basins (Li et al., 2017). High-elevation snowfields are especially important because they store winter precipitation longer and release meltwater later in the season, when water demand is highest. Despite well-documented warming and snow loss, major uncertainties remain regarding how climate change will reshape mountain environments at regional and local scales (Pepin et al., 2025). Recent evidence indicates that snow storage capacity is already declining across western North America, with warming driving earlier melt and altered runoff timing (Hale et al., 2023). Because snow persistence is most sensitive near the freezing threshold, these changes are strongly elevation dependent, with higher elevations currently acting as a relative stronghold for seasonal snow storage and exhibiting slower declines in snow water equivalent than lower-elevation snowpacks (Mote et al., 2005; Hale et al., 2023).

Snow algal blooms have been shown to substantially enhance energy absorption within the snowpack, leading to measurable losses of snow water equivalent even when blooms occupy only a limited fraction of the snow surface, as demonstrated in Western North America (Engstrom et al., 2022; Healy & Khan, 2023). Our results further suggest that biological processes interact with elevational gradients in ways that are hydrologically relevant. If algal-driven darkening is enhanced in upper snowfields through elevation-dependent physiological adjustments, and snow cover is increasingly confined to these elevations, even modest increases in melt efficiency could disproportionately affect the most persistent components of the snowpack, altering the timing of meltwater release. Together, these findings indicate that cellular-scale acclimation of snow algae may influence snowmelt dynamics at broader spatial scales, with implications for water-resource management as warming increasingly affects high-elevation environments.

## Acknowledgements

We thank Katie Coates, Kyle Crichton, Janelle Groff, Andrew Ratz, Victoria Rebbeck, Joshua Vollmer and Jeff Havig for their extensive field and laboratory assistance in mountain snow sample collection and processing. We are grateful to the USDA US Forest Service (Custer-Gallatin, Caribou-Targhee, Bridger-Teton, Flathead, Mt. Baker-Snoqualmie, Gifford Pinchot, Mt. Hood, Deschutes, and Shasta-Trinity National Forests, including the Zigzag Ranger District), the National Park Service (Crater Lake, Glacier, Grand Teton, and Mount Rainier National Parks), and the Natural Resources Department of the Confederated Salish and Kootenai Tribes for sampling permits. We are also grateful to the staff at Mt. Bachelor for facilitating access. We would like to recognize the careful work of the SIF (UC Davis) and the METAL (Arizona State University) in measuring the particulate N component of snow particulates retrieved on filters. This study was funded by the National Science Foundation with grants to TLH (DEB-#2113784) and to JJE (DEB-#2113783).

## Authors contribution

All authors planned and designed the research and conducted fieldwork. JG performed the cell counts. PA conducted all the other analyses and analyzed the data. PA wrote the initial manuscript. All authors contributed to the final version of the manuscript and approved the submitted version.

## Data availability

Sequence Accessions – sequence data are archived at the SequenceRead Archive (SRA) at NCBI under the accession: BioProject (PRJNA1244840).

## Supporting Information

**Supplementary Figure S1.** Spatial distribution of sampling sites across the study regions. Points represent individual samples and are colored by (A) elevation (m a.s.l), (B) day of year (DOY), and (C) snow depth (cm).

**Supplementary Table S1**. Metadata and physicochemical, biological, and pigment-related measurements for all snow samples included in this study. Variables include site and plot identity, sampling date and day of year (DOY), presence of visible snow algae, ecoregion, geographic location, elevation, snow depth, albedo (350–2500 nm), normalized snow water index (NSWI), snow density, cell abundance, particulate carbon (PC) and nitrogen (PN) concentrations, PC:PN ratio, astaxanthin and chlorophyll a content per cell, Astaxanthin:Chl-a ratio, and the fraction of chlorophyll a contained within the upper 2.5 cm of the snowpack. NA indicates data not available.

**Supplementary Table S2**. Top 50 most abundant OTUs across all samples based on total read abundance. Taxonomic assignments were determined using BLAST searches against the NCBI nucleotide database. Query cover and sequence identity correspond to the best BLAST hit for each OTU.

**Supplementary Table S3**. Amplicon sequencing results for all snow samples included in this study. Variables include site, elevation, total algal read counts, algal OTU richness, and the relative abundance (%) of algal taxa based on sequence similarity searches against reference databases. “No blast hit” denotes sequences with <99.4% identity to any reference sequence. NA indicates data not available.

**Supplementary Table S4**. Pearson correlations (r) between elevation and biological, chemical, physical, and optical variables across all samples (Combined) and within each region (Cascade Range and Rocky Mountains). For each variable, sample size (n), number of outliers removed, correlation coefficient (r), and P values are reported. Statistical significance after Bonferroni correction is indicated. Biological variables include cell abundance, biovolume, particulate carbon (PC), nitrogen, PC:PN ratio, and pigment metrics (astaxanthin and chlorophyll-a per cell, and their ratio). Physical variables include NSWI, snow density, and snow depth. Optical variables include absolute reflectance (350–2500 nm), reflectance normalized by cell density, and reflectance relative to clean snow.

**Supplementary Table S5**. Cell volume (µm^3^) of snow algae across samples spanning an elevational gradient. Values are reported as mean ± standard deviation for each sample. Elevation is given in meters above sea level (m a.s.l.).

**Supplementary Table S6**. Linear and mixed-effects model results testing whether bloom developmental stage (astaxanthin per cell and astaxanthin:chlorophyll-a ratio) varies with DOY and snow depth. DOY and snow depth were modeled against region and latitude using linear models, and pigment metrics using mixed-effects models with site as a random intercept. Values shown are β ± SE, t, p, R^2^ (when applicable), and site-level variance (%).

**Supplementary Table S7**. Linear mixed-effects model results for Sanguina-dominated samples, evaluating the effects of biological and environmental predictors on the response variable across all samples (Combined) and within each region (Cascade Range and Rocky Mountains). Models include site (snow patch) as a random intercept. Reported values are fixed-effect estimates (β ± SE) and t values. Random-effect variance (site SD), residual SD, intraclass correlation coefficient (ICC), and sample size (observations/sites) are also shown.

**Supplementary Table S8**. Linear mixed-effects model results including all samples (Sanguina-dominated, other algae-dominated, and no visible algae), evaluating the effects of biological and environmental predictors on the response variable across all samples (Combined) and by region (Cascade Range and Rocky Mountains). Models include site (snow patch) as a random intercept. Reported values are fixed-effect estimates (β ± SE) and t values, along with random-effect variance (site SD), residual SD, ICC, and sample size (observations/sites).

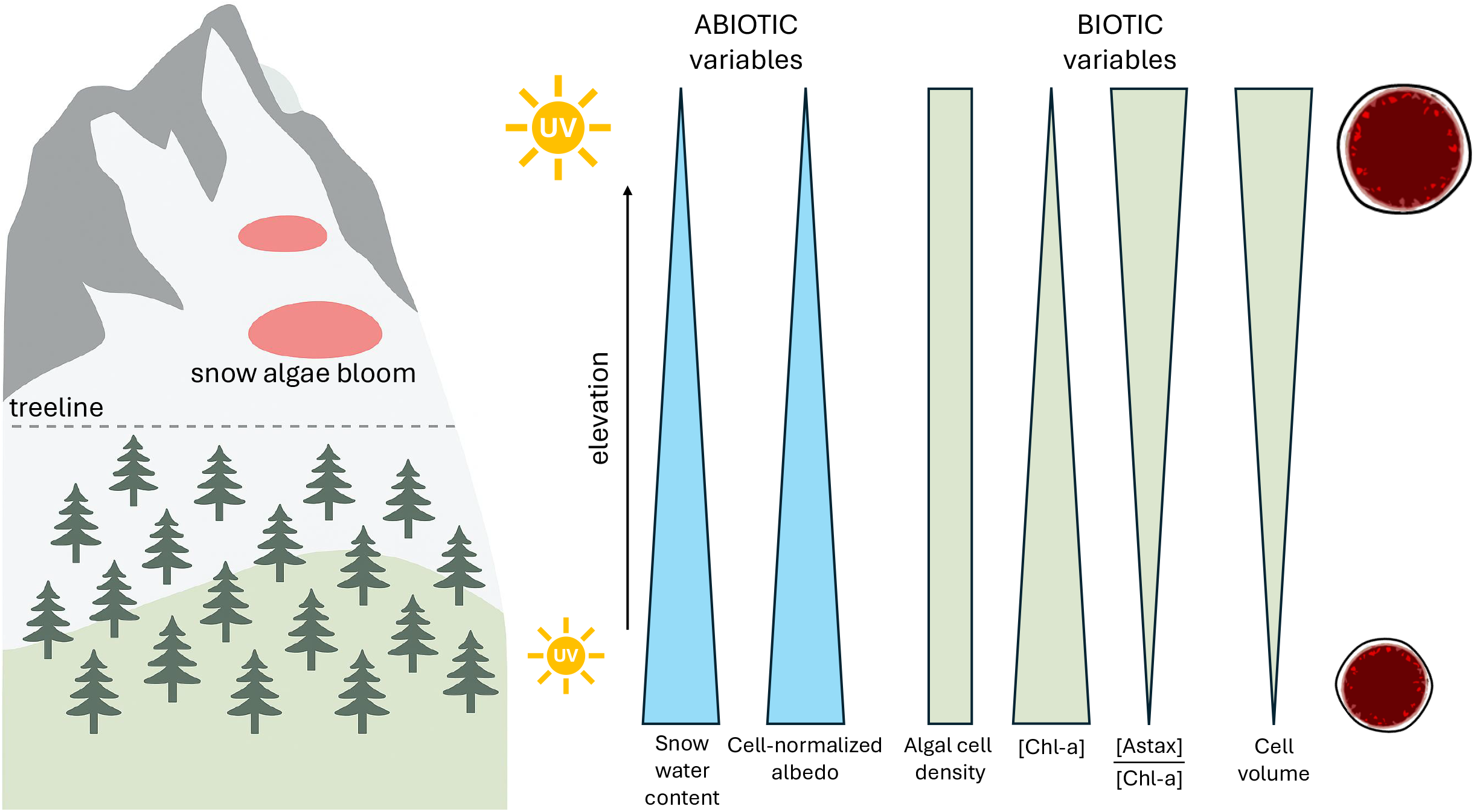

