## Supplementary figures and images for "Elevation shapes alpine snow algal blooms and their influence on albedo reduction"

### Figure S1

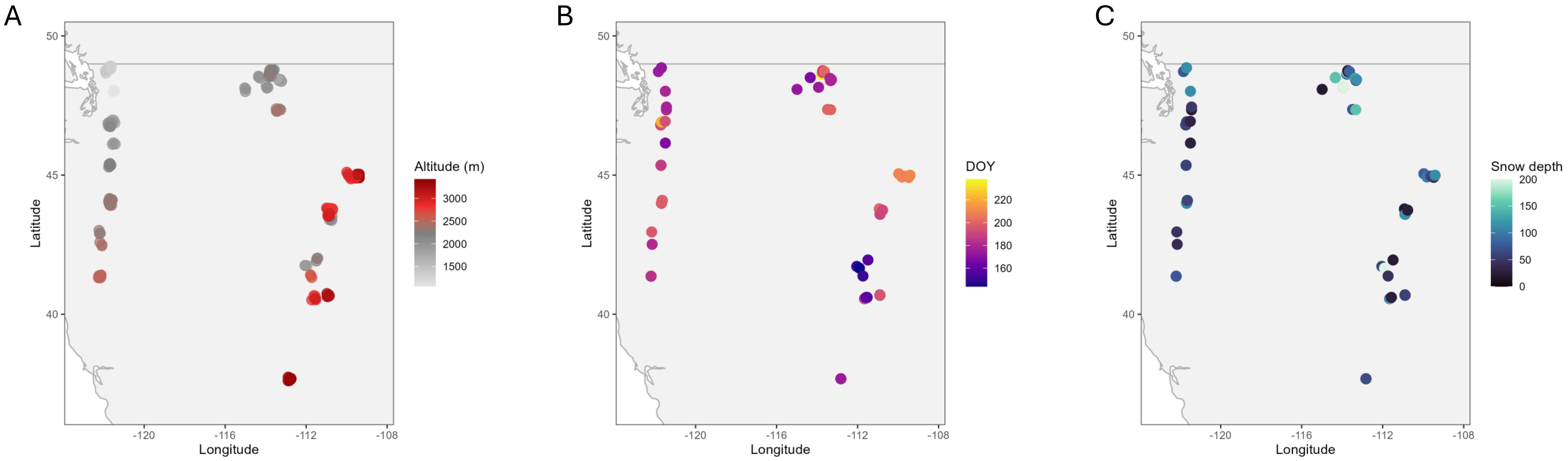
